# Decoding sounds depicting hand-object interactions in primary somatosensory cortex

**DOI:** 10.1101/732669

**Authors:** Kerri M Bailey, Bruno L Giordano, Amanda L Kaas, Fraser W Smith

## Abstract

Neurons, even in earliest sensory regions of cortex, are subject to a great deal of contextual influences from both within and across modality connections. Recent work has shown that primary sensory areas can respond to and in some cases discriminate stimuli not of their target modality: for example, primary somatosensory cortex (SI) discriminates visual images of graspable objects. In the present work, we investigated whether SI would discriminate sounds depicting hand-object interactions (e.g. bouncing a ball). In a rapid event-related functional magnetic resonance imaging (fMRI) experiment, participants listened attentively to sounds from three categories: hand-object interactions, and control categories of pure tones and animal vocalizations, while performing a one-back repetition detection task. Multi-voxel pattern analysis revealed significant decoding of different hand-object interactions within SI, but not for either control category. Crucially, in the hand-sensitive voxels defined from an independent tactile localizer, decoding accuracies were significantly higher for hand-object interactions compared to pure tones in left SI. Our findings indicate that simply hearing sounds depicting familiar hand-object interactions elicit different patterns of activity in SI, despite the complete absence of tactile stimulation. These results highlight the rich information that can be transmitted across sensory modalities even to primary sensory areas.

## Introduction

Much traditional neuroscientific research has investigated the function of the primary sensory brain areas (e.g. visual, somatosensory, and auditory cortices) with respect to how sensory input is processed within its corresponding sensory modality (see Carandini et al. 2005; Kriegeskorte 2015). However, it is well known that primary sensory areas are subject to significant influences from other sensory modalities (for reviews, see Ghazanfar and Schroeder 2006; Driver and Noesselt 2008; see also Lemus et al. 2010; Liang et al. 2013).

Recent studies have used multi-voxel pattern analysis (MVPA) to show that primary sensory cortices can contain fine-grained information about stimuli that are not of their target modality. For example, Meyer et al. (Meyer et al. 2010) showed that simply viewing a silent yet sound-implying video clip triggers discriminable activity in primary auditory cortex (A1) even in the absence of auditory stimulation (see also Vetter et al. 2014 for a complementary example from audition to V1). Recently we have shown that different visual images of familiar but not unfamiliar object categories could be discriminated in primary somatosensory cortex - SI, despite the absence of any tactile stimulation during the experiment (Smith and Goodale 2015; see also Meyer et al. 2011). Thus, there is compelling evidence that stimuli initially presented via one modality can trigger fine-grained activity in distal primary sensory areas for certain pairs of modalities. One possible explanation for these cross-modal effects is that prior experience of the co-occurrence of sensory stimulation across multiple modalities is responsible for such effects (Meyer and Damasio 2009; Smith and Goodale 2015; see also Pérez-Bellido et al. 2018). At the anatomical level there are multiple routes by which information may be transmitted to primary sensory areas that originated from a distal modality: for instance via feedback from higher level multisensory areas (e.g. pSTS, posterior parietal or premotor cortex; see Ghazanfar and Schroeder 2006; Driver and Noesselt 2008), or possibly even via direct connections between specific modalities e.g. from primary auditory to primary visual cortex (e.g. Falchier et al. 2002, 2010).

We note that no study to date has investigated whether fine-grained information about familiar sound categories can be discriminated in SI – despite the fact that anatomical studies in animals suggest direct connections between auditory and somatosensory areas (Cappe and Barone 2005; Ghazanfar and Schroeder 2006) and the demonstration of extensive ipsilateral connections in humans between primary auditory cortex and SI and SII (Ro et al. 2013). As such we would also expect fine-grained auditory responses to be present in early somatosensory areas. Recent work, in fact, has begun to explore whether somatosensory areas may be activated by and discriminate simpler auditory stimuli such as pure tones (Pérez-Bellido et al. 2018). These authors have shown representations of auditory frequency in early somatosensory areas (pooled across likely SI and SII) that relate to perception. The authors propose that these representations of auditory information in somatosensory areas may be related to predictive processing of associated tactile features (see e.g. Friston 2009; Friston et al. 2009; Clark 2013). Zhou and Fuster (Zhou and Fuster 2004), moreover, have shown that neurons in SI respond to auditory cues when they are predictive of upcoming somatosensory stimulation while Lemus et al. (Lemus et al. 2010) showed auditory responses in SI and SII to acoustic flutter stimuli that could not, however, discriminate the stimulus identity presented. In addition, previous work has indicated the activation of somatosensory areas in audio-haptic matching tasks (postcentral gyrus: Kassuba et al. 2013), in listening to tool sounds (Lewis et al. 2005; likely SII; see also Lewis et al. 2006) and in particular sub-regions of SII (OP1 & OP4) while listening to various sound categories (Beauchamp and Ro 2008).

Building on these studies and our earlier work (Smith and Goodale 2015), in the present experiment we set out to investigate whether primary somatosensory cortex (SI) would contain fine-grained information that discriminates sounds depicting familiar hand-object interactions (e.g. bouncing a ball vs typing on a keyboard). We reasoned that such effects would occur in SI due to the prior experience of the co-occurrence of the haptic information that corresponds with hearing the sound of specific hand-object interaction categories (e.g. bouncing a ball vs typing on a keyboard – see Meyer and Damasio 2009). Based on previous univariate studies examining the responses observed to tools and animal sounds, we might also expect sounds depicting hand-actions to preferentially activate (see Lewis et al. 2005) and possibly be discriminated within higher level somatosensory (likely SII) and also premotor areas (via the auditory dorsal stream: Rauschecker and Tian 2000; Bizley and Cohen 2013). While premotor areas are not the focus of the present study, we note that mirroring accounts would also predict such sounds to be discriminated in premotor cortex – most likely in ventral premotor areas (see e.g. Gazzola et al. 2006).

Thus in the present study, participants listened to sound clips of familiar hand-object interactions (e.g. bouncing a ball, typing on a keyboard), in addition to two control categories (animal vocalizations – familiar high level control; pure tones – unfamiliar low level control), in an event-related functional magnetic resonance imaging (fMRI) experiment. Based on our previous work (Smith and Goodale 2015) and the co-occurrence account (Meyer and Damasio 2009) we predicted that MVPA would show significant decoding of sound identity for hand-object interaction sounds in SI, particularly in independently localized tactile hand-sensitive voxels, but not for the two control categories. We further predicted decoding would be greater in SI for hand-object interaction sounds than for the two control categories. We repeated these analyses in a series of additional control regions - including secondary somatosensory cortex and premotor cortex – where multiple accounts may predict reliable decoding for hand-object interaction sounds but possibly also for each additional sound category – as these areas have both been implicated in the dorsal auditory ‘how’ pathway. In early auditory cortex we would expect reliable decoding of all sound categories (positive control), whereas in M1 (see Smith and Goodale 2015; see also Lewis et al. 2005) we did not expect to find any reliable decoding (negative control). We did not have a specific prediction regarding whether decoding would be present in V1 as previous studies finding such effects (Vetter et al. 2014; Gu et al. 2020) have used rich natural soundscapes rather than particular object categories in isolation but we include this region for comparability to previous studies.

## Methods

### Participants

Self-reported right-handed healthy participants (*N* = 10; 3 male), with an age range of 18-25 years (*M* = 22.7, *SD* = 2.41), participated in this experiment. We based our sample size on prior work in this area – investigating cross-sensory effects in early sensory areas with MVPA - that has all used a similar sample size of 10 or fewer (Meyer et al. 2011; Vetter et al. 2014; Smith and Goodale 2015; Pérez-Bellido et al. 2018; see also Wang et al. 2021; de Borst and de Gelder 2017). No power analysis was conducted. All participants reported normal or corrected-to-normal vision, and normal hearing, and were deemed eligible after meeting MRI screening criteria, approved by the Scannexus MRI centre in Maastricht. Written consent was obtained in accordance with approval from the Research Ethics Committee of the School of Psychology at the University of East Anglia. Participants received €24 euros (equivalent to £20 sterling British pounds) for their time.

### Stimuli & Design

Three different categories of auditory stimuli were used in a rapid event-related fMRI design; sounds depicting hand-object interactions, animal vocalizations, and pure tones. There were five different sub-categories within each of these categories, with two exemplars of each sub-category, thus resulting in 30 individual stimuli in total. The five hand-object interaction stimuli consisted of bouncing a basketball, knocking on a door, typing on a keyboard, crushing paper, and sawing wood. These stimuli were chosen for the reason that participants should have previously either directly experienced rich haptic interactions with such objects, or observed such interactions. Two control categories were also used. Firstly, animal vocalizations were used as familiar sounds not directly involving interactions with the hands. These consisted of birds chirping, a dog barking, a fly buzzing, a frog croaking, and a rooster crowing. An independent ratings experiment confirmed these sounds were matched to the hand-object interactions for familiarity (1-7 scale: *M* = 5.31 for both hand-object interactions and animal vocalizations; see Supp. Table 1) and a further similarity rating experiment with an independent group of participants confirmed that the exemplars within the familiar sound categories were also matched for within-category similarity (see Supp. Figure 1 and Supp. Text). Sounds from these two categories were downloaded from SoundSnap.com, YouTube.com, and a sound database taken from Giordano, McDonnell, and McAdams (Giordano et al. 2010). The second control category were non-meaningful sounds, defined as pure tones. These consisted of pure tones of five different frequencies (400Hz, 800Hz, 1600Hz, 3200Hz, and 6400Hz; 2000ms duration), created in MatLab. All sounds were stored in WAV format, and were cut to exactly 2000ms using Audacity 2.1.2, with sound filling the entire duration. Finally, all sounds were normalised to the root mean square level (RMS: Giordano et al. 2013). More information regarding how these sounds were selected, including pilot experiments and ratings for the sounds, can be seen in Supp. Text and Supp. Table 1.

Each run began and ended with 12s silent blocks of fixation. After the initial 12s fixation, 60 individual stimuli were played, with each unique sound presented twice per run. Stimuli were played in a pseudo-randomly allocated order at 2s duration with a 3s ISI (5s trial duration; no jitter). At random intervals, 15 null trials (duration 5s) were interspersed where no sound was played. This resulted in a total run time of 399s. We note that rapid event related designs are often used in perception research with relatively short trial lengths and hence ISIs (see e.g. Giordano et al 2021; see also Smith & Muckli 2010; Petro et al 2013) as a way of maximizing the number of repetitions obtained. We used small trial lengths with more trials in order to have more data for training and testing the MVPA classifier, which is an important consideration (see e.g. Pereira et al 2009). After the main experiment, a somatosensory localiser was included to map the hand region in the somatosensory cortex (see Smith and Goodale 2015). Piezo-electric Stimulator pads (Dancer Design, UK) were placed against the participant’s index finger, ring finger, and palm of each hand using Velcro (six pads total, three per hand; see Supp. Figure 8 for a visual example on one hand). Each pad contained a 6mm diameter disk centred in an 8 mm diameter static aperture. The disks stimulated both hands simultaneously with a 25Hz vibration in a direction normal to the surface of the disk and skin, at an amplitude within the range of ±0.5mm). Localiser runs consisted of 15 stimulation blocks and 15 baseline blocks (block design, 12s on, 12s off, 366s total run time). Note that for the first two participants, a slightly modified timing was employed (block design, 30s on, 30s off; 10 stimulation blocks, 9 baseline blocks).

### Procedure

After signing informed consent, each participant was trained on the experimental procedure on a trial set of stimuli not included in the main experiment, before entering the scan room. Participants were instructed to fixate on a black and white central fixation cross presented against a grey background whilst listening carefully to the sounds, which were played at a self-reported comfortable volume (as in Leaver and Rauschecker 2010; Meyer et al. 2010; Man et al. 2012, 2015). Participants performed a one-back repetition counting task, and hence counted the number of times they heard any particular sound repeated twice in a row, for example, two sounds each of a dog barking (randomly allocated from 2 to 6 per run). We chose this task as it was important that the task did not require an explicit motor action such as pressing a button, to prevent a possible confound in somatosensory cortex activity (see Smith and Goodale 2015). Thus, participants verbally stated the number of counted repetitions they heard at the end of each run, and they were explicitly asked to not make any movements in the scanner unless necessary. Overall, most participants completed either 8 or 9 runs (with the exception of one participant, who completed 7), thus, participants were exposed to approximately 32-36 repetitions per stimulus, and 16-18 repetitions per unique sound exemplar. After the main experiment, participants took part in the somatosensory mapping localiser. Participants were not informed about this part of the experiment until all main experimental runs had been completed. Each participant completed 1 (*N* = 2) or 2 (*N* = 8) somatosensory mapping runs, and kept their eyes fixated on a black and white central fixation cross presented against a grey background for the duration of each run. Participants were debriefed after completion of all scanning sessions.

### MRI Data Acquisition

Structural and functional MRI data was collected using a high-field 3-Tesla MR scanner (Siemens Prisma, 64 channel head coil, Scannexus, Maastricht, The Netherlands). High resolution T1 weighted anatomical images of the entire brain were obtained with a three-dimensional magnetization-prepared rapid-acquisition gradient echo (3D MPRAGE) sequence (192 volumes, 1mm isotropic). Blood-oxygen level dependent (BOLD) signals were recorded using a multiband echo-planar imaging (EPI) sequence: (400 volumes, TR = 1000ms; TE = 30ms; flip angle 77; 36 oblique slices, matrix 78 × 78; voxel size = 2.5mm^3^; slice thickness 2.5mm; interslice gap 2.5mm; field of view 196; multiband factor 2). A short five volume posterior-anterior opposite phase encoding direction scan was acquired before the main functional scans, to allow for subsequent EPI distortion correction (Jezzard and Balaban 1995; Fritz et al. 2014). Slices were positioned to cover somatosensory, auditory, visual, and frontal cortex. Sounds were presented via an in-ear hi-fi audio system (Sensimetrics, Woburn MA, USA), and the visual display was rear projected onto a screen behind the participant via an LCD projector. Finally, a miniature Piezo Tactile Stimulator (mini-PTS; developed by Dancer Design, UK) was used to deliver vibro-tactile stimulation to the hands, using the same fMRI sequence with a modified number of volumes (366s for the majority, slightly longer for the first two participants due to slightly different design – see Stimuli & Design above).

### MRI Data Pre-processing

Functional data for each main experimental run, in addition to somatosensory localiser runs, was pre-processed in Brain Voyager 20.4 (Brain Innovation, Maastricht, The Netherlands; Goebel et al. 2006), using defaults for slice scan time correction, 3D rigid body motion correction, and temporal filtering. Functional data were intra-session aligned to the pre-processed functional run closest to the anatomical scan of each participant. Distortion correction was applied using COPE 1.0 (Fritz et al. 2014), using the 5 volume scan acquired in the opposite phase encode direction (posterior to anterior) for each participant. Voxel displacement maps (VDM)’s were created for each participant, which were applied for EPI distortion correction to each run in turn. Functional data were then coregistered to the participant’s ACPC anatomical scan. Note no Talairach transformations or spatial smoothing were applied, since such a transformation would remove valuable fine-grained pattern information from the data that may be useful for MVPA analysis (Fischl et al. 1999; Argall et al. 2006; Goebel et al. 2006; Kriegeskorte and Bandettini 2007). For the main MVPA analyses (described further below) we conducted a GLM analysis independently per run per participant, with a different predictor coding stimulus onset for each unique stimulus presentation (3 categories X 5 within-sub categories X 2 unique exemplars, each repeated twice = 60 predictors) convolved with a standard double gamma model of the haemodynamic response function (see Smith and Muckli 2010; Greening et al. 2018). The resulting beta-weight estimates are the input to the pattern classification algorithm described below (see Multi-Voxel Pattern Analysis).

The somatosensory mapping localiser data was analysed using a GLM approach, with one predictor defining stimulation onset convolved with the standard double gamma model of the haemodynamic response function. The *t*-values were defined from the somatosensory localiser by taking the contrast of stimulation vs baseline in each participant. This allowed us to define the 100 voxels showing the strongest tactile response to hand stimulation in each individual’s hand-drawn anatomical mask (see Regions of Interest below) of the post central gyrus (see Smith and Goodale 2015; Perez-Bellido et al 2018; for comparable analyses).

### Regions of Interest

#### Anatomical mask of SI

In order to accurately capture the potential contribution from each sub-region of SI – i.e. area 3a, 3b, 1 or 2, hand-drawn masks of the PCG were created in each individual participant (Meyer et al. 2011; Smith and Goodale 2015). This allowed us to go beyond the capabilities of the somatosensory localiser alone, by enabling inclusion of all the information potentially available – i.e. both tactile and proprioceptive - in SI for the pattern classification algorithms (see Smith and Goodale 2015 for further information).

Anatomical masks were created using MRIcron 6 (Rorden et al. 2007) using each participant’s anatomical MRI scan in ACPC space. As in Meyer et al. (Meyer et al. 2011) and Smith and Goodale (Smith and Goodale 2015), the latero-inferior border was taken to be the last axial slice where the corpus callosum was not visible. From anterior to posterior the masks were defined by the floors of the central and post-central sulci. Furthermore, masks did not extend to the medial wall in either hemisphere (Meyer et al. 2011; Smith and Goodale 2015). This resulted in an average of 41 slices (total range 39 to 46) for each hemisphere per participant. The average voxel count was 1969 (*SD* = 229) for the right PCG, and 2106 (*SD* = 215) for the left PCG, which did not significantly differ from one another (see Figure 1 and Supp. Figures 2 & 3).

**Figure 1:**
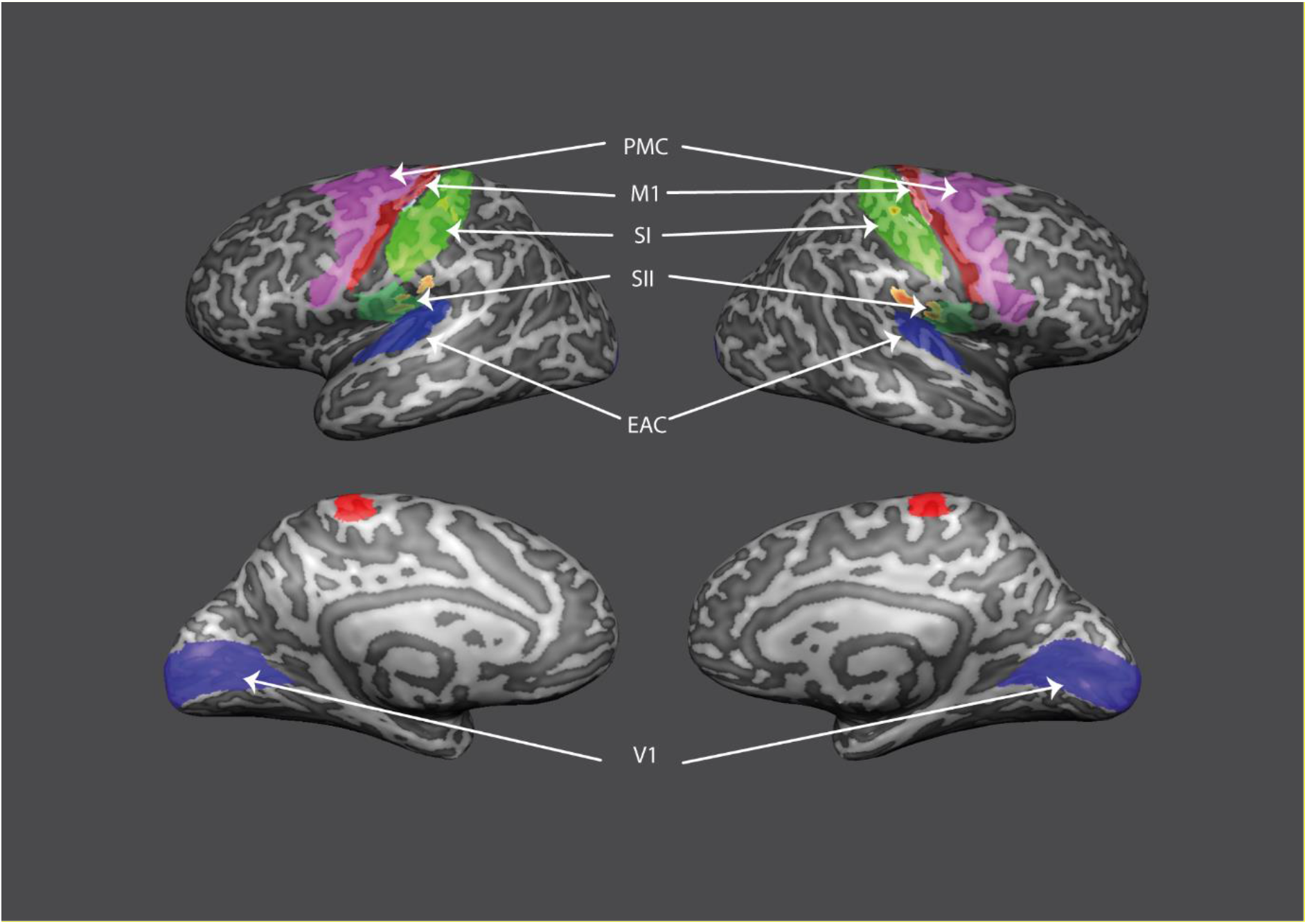
Hand drawn anatomical mask of the SI ROI (Green) in a representative participant shown on an inflated cortical reconstruction of each hemisphere and in both medial and lateral views. Additional ROIs defined from the Glasser parcellation are also shown overlaid. The underlying activation shows the stimulation vs baseline contrast from the somatosensory hand localizer for the same participant (*t* > 3, *p* = .002, uncorrected). Warm colours signify stimulation > baseline and cool colours stimulation < baseline.

#### SI Tactile Mask

We also created a sub-sampled version of the SI anatomical mask per participant which comprised the 100 most activated voxels in response to tactile stimulation of the hands from the tactile localizer scan (see above). The rationale for including this additional ROI is two-fold: first, it allows us to isolate whether any effects are present within tactile sensitive voxels in S1 as has been evident in our previous work (see Smith & Goodale, 2015; see also Perez-Bellido et al 2018 for a comparable approach). Second, the analysis permits the use of a much small number of voxels (100 vs ~2000) allowing the influence of ROI size to be taken into account. An example of the somatosensory localizer activation is shown in Figure 1 (see Supp. Figure 4 for group activation). From this point onwards, this ROI will be referred to as SI - Tactile.

#### Additional Regions of Interest

In order to define a series of additional ROIs we used Freesurfer (version 6.0.0) together with the Glasser parcellation (Glasser et al. 2016). The atlas annotation files were converted to fsaverage coordinates (Fischl et al. 1999) and from there mapped to each individual participant’s cortical reconstruction via spherical averaging (see Kietzmann et al. 2019). We defined each region, independently per hemisphere, according to the detailed specification given in Glasser et al. (Glasser et al. 2016): Early Auditory Cortex (Areas M, P & L Belt, A1 and RI), Pre-Motor cortex (Areas 6a, 6d, 6r, 6v, FEF, PEF and area 55b), Primary Motor Cortex (Area 4), Secondary Somatosensory Cortex (Areas OP1-4; see Eickhoff, Amunts, et al. 2006; Eickhoff, Schleicher, et al. 2006; Keysers et al. 2010) and Primary Visual Cortex (Area V1). Finally, we used the co-registration procedure in Brain Voyager to align the native space anatomy from Free Surfer with that of the functional data in ACPC space. In Figure 1 we show each of these regions in surface space for a representative participant – and in Supp. Figure 3 these regions are displayed for each individual participant (see Supp. Table 2 for mean volume for each additional ROI).

#### Multi-Voxel Pattern Analysis

For the multi-voxel pattern analysis (MVPA; e.g. Haynes 2015), we trained a linear support vector machine (SVM) to learn the mapping between the spatial patterns of brain activation generated in response to each of the five different sounds within a particular sound category (for example: for hand-object interactions, the classifier was trained on a five way discrimination between each relevant sub-category: typing on a keyboard, knocking on a door, crushing paper and so on; Smith and Muckli 2010; Vetter et al. 2014; Smith and Goodale 2015; Greening et al. 2018). The classifier was trained and tested on independent data, using a leave one run out cross-validation procedure (Smith and Muckli 2010; Smith and Goodale 2015). The input to the classifier was always single trial brain activity patterns (beta weights) from a particular ROI while the independent test data consisted of an average activity pattern taken across the repetitions of specific exemplar in the left out run (e.g. the single trial beta weights of the four presentations of ‘bouncing a ball’ in the left out run were averaged). We have used this approach successfully in previous studies, as averaging effectively increases the signal to noise of the patterns (Smith and Muckli 2010; Vetter et al. 2014; Muckli et al. 2015). For similar approaches applied to EEG and MEG data, see Smith and Smith (Smith and Smith 2019) and Grootswagers, Wardle, and Carlson (Grootswagers et al. 2017) respectively.

Finally, we used the LIBSVM toolbox (Chang and Lin 2011) to implement the linear SVM algorithm, using default parameters (*C* = 1). LIBSVM uses the one vs one method for multiclass classification (Chang and Lin 2011). The activity pattern estimates (beta weights) within each voxel in the training data was normalised within a range of −1 to 1, prior to input to the SVM (Smith and Muckli 2010; Chang and Lin 2011; Vetter et al. 2014; Smith and Goodale 2015; Greening et al. 2018; Knights et al. 2021). The test data were also normalised using the same parameters as in the training set, in order to optimise classification performance. To test whether group level decoding accuracy was significantly above chance, we performed non-parametric Wilcoxon signed-rank tests using exact method on all MVPA analyses, against the computed empirical chance level (Formisano et al. 2008; Greening et al. 2018), with all significance values reported two-tailed. We used a permutation approach – randomly permuting the mapping between each condition and each label, independently per run, to calculate the empirical chance level for each participant and each decoding analysis separately (we note here that the average empirical chance level across all participants, regions and analyses was 0.202, with a standard deviation of .0061, which is very close to the theoretical chance level of 0.20). We implemented control for multiple comparisons by using the False Discovery Rate (q < .05) method (Benjamini and Yekutieli 2001). Effect sizes are reported for the Wilcoxon tests using the Z/sqrt(N) formula (Rosenthal 1991). In each region where decoding accuracy was significant for any category, we conducted follow-up tests between each sound category again with Wilcoxon signed rank tests (two tailed, FDR corrected for multiple comparisons across all regions showing significant decoding effects; see Knights et al. 2021)

#### Univariate Deconvolution Analysis

In order to examine whether there were univariate differences between sound categories in each ROI we performed a deconvolution analysis (see Smith and Muckli 2010; Petro et al. 2014). In this analysis, there were 20 predictors (each TR=1s here) to cover the full extent of the HRF response for each stimulus presented in a given condition (so 60 predictors in total, plus confounds for each experimental run). We then used an ANOVA approach to ask whether the average beta weight across time points 5-8s, corresponding to the peak of the HRF, were significantly different across sound categories, hemisphere and ROI.

#### Erosion Analysis

We also performed a subsequent decoding analysis where we eroded the closest voxels between neighbouring (EAC and SII) and relatively near (SI and PMC) ROIs that displayed significant effects in a systematic manner in order to demonstrate that the decoding effects found in nearby areas are not trivially due to partial volume effects or other spatial artifacts (Pérez-Bellido et al. 2018). Our approach (see Pérez-Bellido et al. 2018) involved computing the Euclidean distance between each voxel’s position in each relevant ROI pair (in ACPC space) and then removing those voxels deemed closest in several discrete steps (1-7mm; see Supp. Figure 6). We show FDR-corrected decoding accuracy as a function of distance to the nearest voxel in the neighbouring ROI (see Figure 3). We also plotted the univariate signal as a function of distance to the nearest voxel (see Supp. Figure 7).

## Results

Bilateral anatomical masks of the lateral postcentral gyri (SI) were manually defined in each participant (see Figure 1 & Methods) whilst a subject-specific cortical parcellation was used to define a set of additional regions of interest (see Methods). We computed cross-validated decoding performance of sound identity for each sound category independently (hand-object interactions; animal vocalizations, and pure tones control) in each ROI, and report FDR-corrected effects on decoding accuracy below.

### Primary Somatosensory Cortex

As expected, significantly above-chance decoding was found for hand-object interaction sounds in left SI (Med = 30.56%; *p* = .0195; *d* =.73), right SI (Med = 23.65%; *p* = .0273; *d* = .69) and pooled SI (Med = 28.75%; *p* = .0098; *d* = .79); signed rank, two-tailed test, versus subject-specific empirical chance level, FDR corrected (see Figure 2A). Crucially, the same analyses for our two control categories of familiar animal vocalizations and unfamiliar pure tones did not show any significant decoding above chance in right, left, or pooled SI (all *p’s* > .432). When pooled across hemispheres, SI showed reliably greater decoding for hand-object interactions than pure tones (*p* = .0156, *d* = .75; FDR corrected, signed rank, two-tailed test). No other differences between sound categories in any sub-region were significant (all *p’s* > .0527). The confusion matrices underlying the classification performance can be seen in Supp. Figure 4. Thus, SI carries content-specific information for the hand-object interaction sounds which convey haptic properties with the hands, regardless of hemisphere.

**Figure 2:**
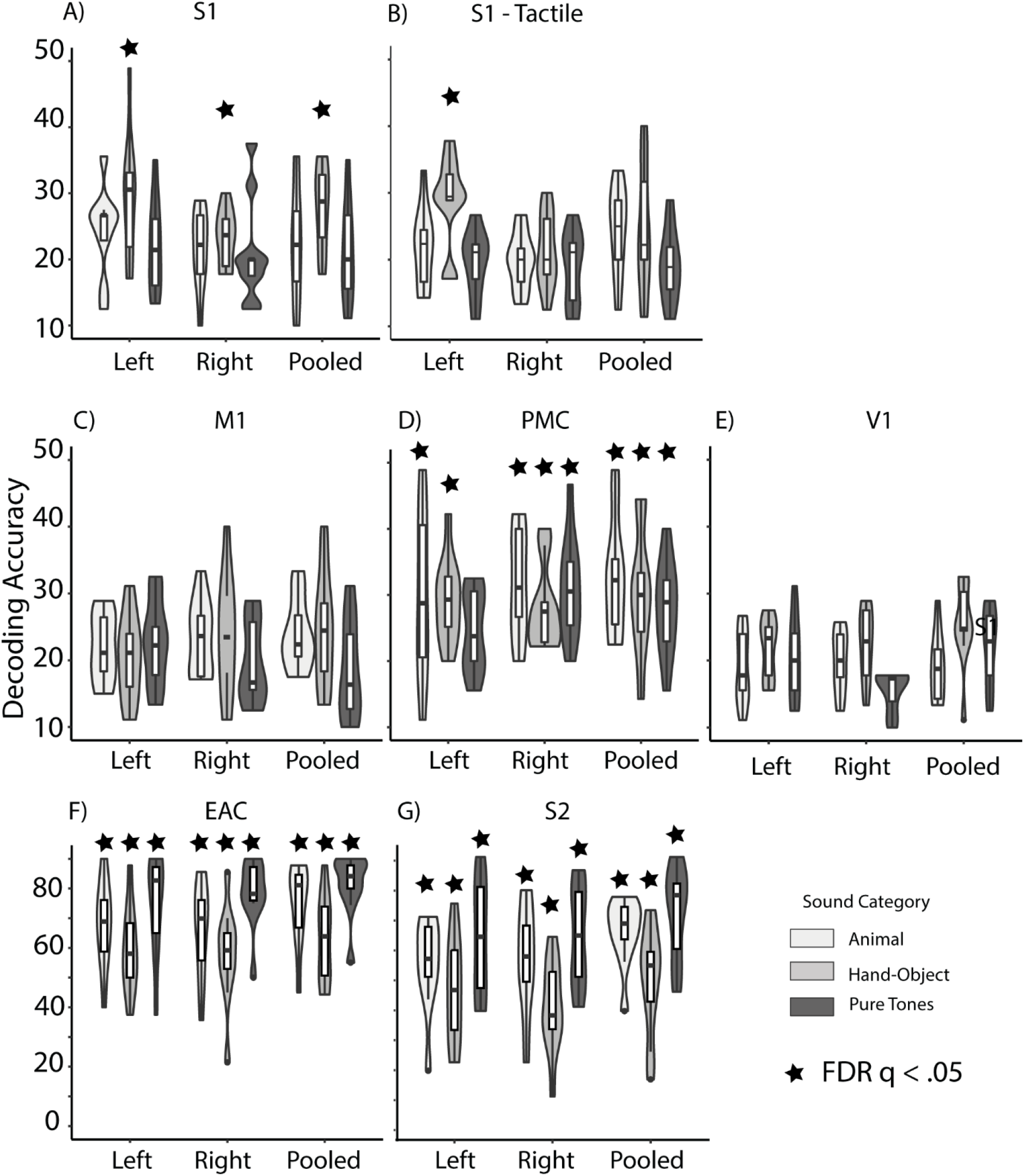
Decoding of sound identity. (A) Violin plot depicting cross-validated 5AFC decoding performance for each stimulus category (animal vocalizations, hand-object interactions and pure tones) for the right and left SI independently and pooled across hemispheres. Violin plot shows the full range of the underlying data while the boxplot depicts the median and the IQ range. * = significant after FDR correction. (B) As in A but for the top 100 voxels that were responsive to tactile stimulation of the hands in an independent localizer session. (C–F) As in A but for several additional regions of interest.

**Figure 3:**
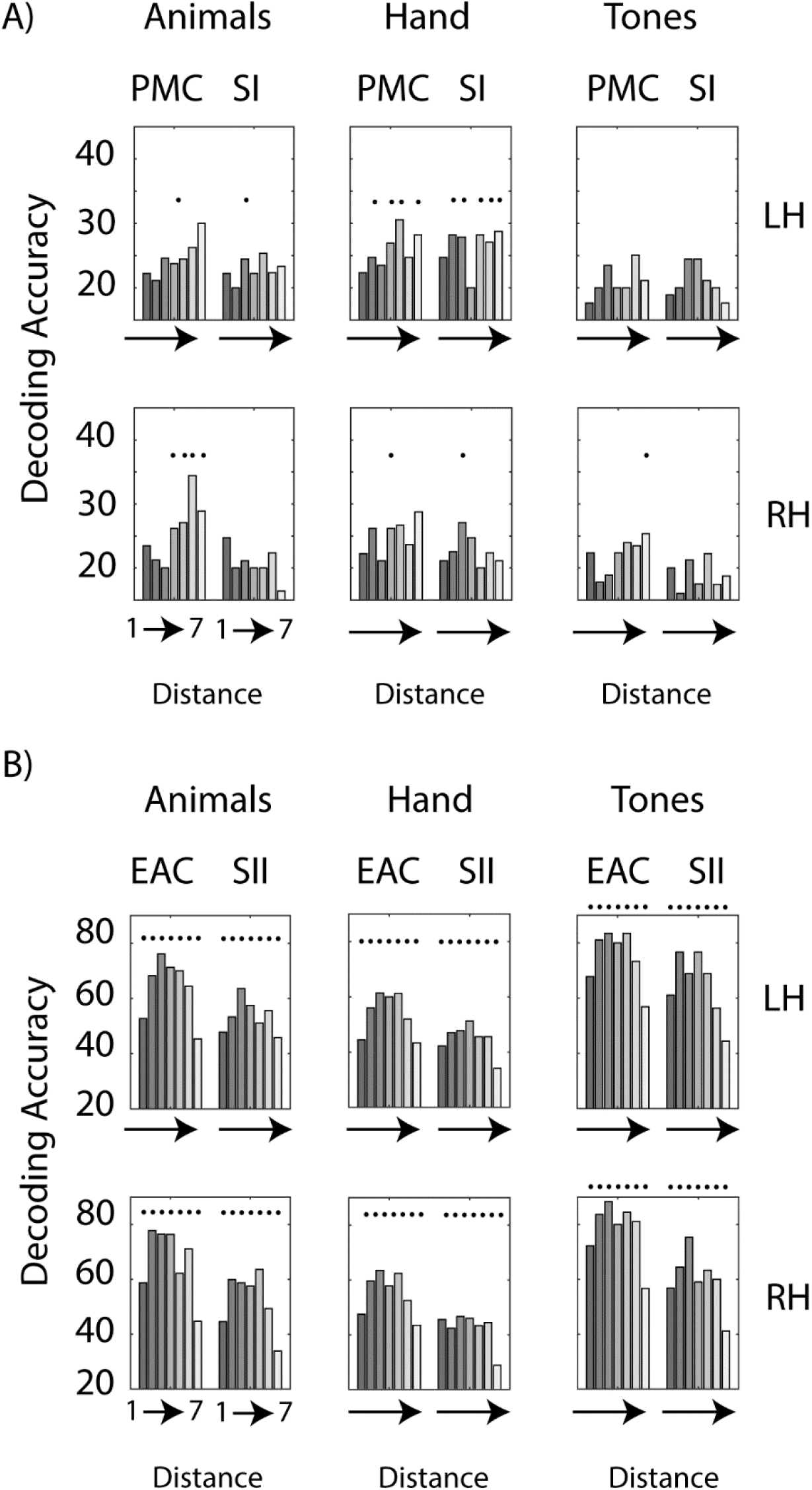
Erosion Analysis. (A) Decoding of sound identity in PMC and SI for each sound category and each ROI (split by hemisphere), as a function of the level of voxel erosion employed to eliminate the contribution of nearby voxels (1-7mm). (B) As in A except for EAC and SII. * = significant after FDR correction.

We sought to corroborate the above finding when selecting the top 100 most active voxels in each sub-region of SI from the independent somatosensory localiser (SI – Tactile; see Methods) as an independent means of selecting a subset of sensitive voxels in the present task (see Smith and Goodale 2015; see also Pérez-Bellido et al. 2018). These analyses revealed significant decoding for hand-object interactions only in left SI (Med = 29.44%; *p* = .009, *d* = .79); signed rank, two-tailed test, versus subject-specific empirical chance level, FDR corrected (see Figure 2B). Critically, decoding accuracies for hand-object interactions in left SI were significantly higher than for pure tones (Hands vs Tones: *p* = .006, *d* = .82) while the predicted difference between hand-object interactions and animal vocalizations was not significant after FDR correction (Hands vs Animal Vocalizations: *p* = .0195, *d* =.55). In agreement with the previous analyses using the entire SI, no other sound categories were decoded above chance (all *p’s* ≥ .232). These results show the classifier could reliably decode only hand-object interaction sounds above chance when constrained to the hand-sensitive voxels in left SI, with accuracies that were significantly higher than the pure tone control sounds. Thus sounds depicting hand-object interactions were reliably discriminated when restricting the MVPA analysis to voxels with high responses to tactile stimulation of the right, but not left, hand.

### Additional Regions of Interest

Using a recently published cortical surface based parcellation atlas (Glasser et al. 2016) we defined several additional control regions that we expected to be important in the present task (see Figure 1 & Methods; see also Smith and Goodale 2015).

#### Early Auditory Cortex

As would be expected, decoding in early auditory cortex (EAC) was robustly significant for all sound categories (all Meds ≥ 68%, all *p*’s ≤ .002, all *d’s* > =.884; signed rank, two-tailed test, versus subject-specific empirical chance level, FDR corrected: see Figure 2C). In each sub-region of EAC decoding accuracy was significantly better for pure tones than hand object interactions (all *p’s* ≤ .0059, all *d’s* ≥ .82), whilst pure tones were also decoded better than animal vocalizations in Left and Right EAC (both *p*’s ≤ .0078, both *d’s* ≥ .81). Furthermore, decoding was greater for animal vocalizations than hand-object interactions in both Left and Right EAC (both *p*’s ≤ .0117, both *d’s* ≥ .77). Thus in EAC, all sound categories were highly discriminated with the specific pattern of decoding suggesting the opposite to that in SI, with better decoding of pure tones, followed by animal vocalizations, then hand-object interaction sounds.

#### Secondary Somatosensory Cortex

Decoding in SII was robustly significant for all sound categories in each hemisphere (all Meds ≥ 47.80%, all *p*’s ≤ .0039, all *d’s* ≥ .85; signed rank two-tailed test versus subject-specific empirical chance level, FDR corrected; see Figure 2C). In Left SII, decoding accuracy was significantly better for pure tones than for hand-object interactions (*p* = .004, *d* = .85) whilst the difference between pure tones and animal vocalizations was not significant (*p* = .080, *d* = .56). In Right SII, decoding was significantly higher for animal vocalizations than hand-object interactions (*p* = .004, *d* = .85), and for pure tones than for hand-object interactions (*p* = .002, *d* = .89) but again there was no significant difference between animal vocalizations and pure tones (*p* = .078, *d* = .58). When pooling across hemispheres, higher decoding for animal vocalizations than hand-object interactions (*p* = .002, *d* = .89) and for pure tones than hand-object interaction (*p* = .004, *d* = .84) was found. Thus in SII better decoding was found for both pure tones and animal vocalizations than for hand-object interactions.

#### Pre-Motor Cortex

In pre-motor cortex (PMC), significantly above chance decoding was found for hand-object interactions in right PMC (Med = 27.5%; *p* = .002, *d* = .89), left PMC (Med = 29.31%, *p* = .0039, *d* = .85) and pooled PMC (Med = 31.25%, *p* = .0098, *d* = .79), signed rank two-tailed test versus subject-specific empirical chance level, FDR corrected (see Figure 2D). In addition, significant decoding of animal vocalizations was found in Left (Med = 28.80; *p* = .0273, *d* = .69), Right (Med = 33.1; *p* = .004, *d* = .85) and Pooled PMC (Med = 32.2; *p* = .002, *d* = .89), and of pure tones again in Right (Med = 30.54; *p* = .004, *d* = .86) and Pooled PMC (Med = 28.89; *p* = .002, *d* = .79). There were no significant differences in decoding accuracy after FDR correction in any ROI (all *p’s* > .107, all *d’s* ≤ .51). Thus PMC contains information about multiple types of sound category in each hemisphere.

#### Primary Motor Cortex

Decoding accuracies in primary motor cortex (M1) revealed no reliable above-chance decoding after FDR correction (all *p’s* > 0.13, all *d’s* ≤ .50; signed rank two-tailed test versus subject-specific empirical chance level, FDR corrected; see Figure 2E).

#### Primary Visual Cortex

Decoding accuracies in primary visual cortex (V1) revealed no significant above chance decoding after FDR correction (all *p’s* ≥ .037, all *d’s* ≤ .66; signed rank two-tailed test versus subject-specific empirical chance level, FDR corrected; see Figure 2F).

### Erosion Analysis

As some regions showing significant decoding are either directly neighbouring to one another (EAC and SII) or relatively close at specific locations (SI and PMC) and showed similar decoding effects, we performed a control analysis (Figure 3) where we eroded a portion of the voxels (in 1mm steps from 1-7 mm, see Methods & Supp. Fig 5) from each ROI to minimize the contribution of partial volume effects or other spatial artifacts to the effects observed (see Pérez-Bellido et al. 2018). These analyses reveal that decoding remained significant even when the closest 3mm voxels were discarded in both Left SI and Left PMC for hand-object interaction sounds (two tailed signed rank tests against subject-specific empirically determined chance levels, FDR corrected - see Figure 3A). In addition, significant effects were present in each region of EAC and SII for each sound category at all eroded voxel levels (two tailed signed rank tests against per participant empirically determined chance levels, FDR corrected, see Figure 3B). Thus the key effects observed in somatosensory brain areas to sound stimuli are unlikely to be due to partial volume or spill-over effects from neighbouring or nearby regions that also display significant effects in the present experiment.

### Univariate Analysis

Finally, we examined the univariate responses to each stimulus category in each ROI (Figure 4: deconvolution analysis). As expected, strong responses were evident in EAC but also in SII. However, the signal was much weaker in SI, M1, PMC and V1. A three-way ANOVA (Sound Category, Hemisphere and Region) revealed a significant three-way interaction (*F*_20, 180_ = 4.60, *p* < .001), as such separate 2-way ANOVAS were run in each region separately. There were no reliable main effects or interactions in SI, V1, M1 (all *p*’s > .179) or in PMC (all *p*’s > .053). However in EAC there was a significant interaction between Sound Category and Hemisphere: *F*_2.79, 25.15_ = 7.25, *p* = .001. In each sub-region of EAC however, both animal vocalizations and hand-object interactions produced stronger activity than pure tones (all corrected *p*’s ≤ .0167) but there was no difference between the two former categories (both *p’s* ≥ .639). In contrast, in SII only the effect of Sound Category was significant (*F*_1.60, 14.40_ = 9.73, *p* =.003), with higher amplitude for animal vocalizations than either hand-object interactions or pure tones (corrected *p’s* ≤ .016) but no significant difference between the latter two (corrected *p* = .099). Thus there is preferential activation of EAC for familiar sound categories (hand object interactions and animal vocalizations) versus pure tones, whereas in SII preferential activation is observed only for the animal vocalizations versus each other category. In SI however, there is no evidence of significant univariate differences between sound categories (we note that the same is true for the tactile-sensitive voxel subset – all *p’s* ≥ .33).

**Figure 4:**
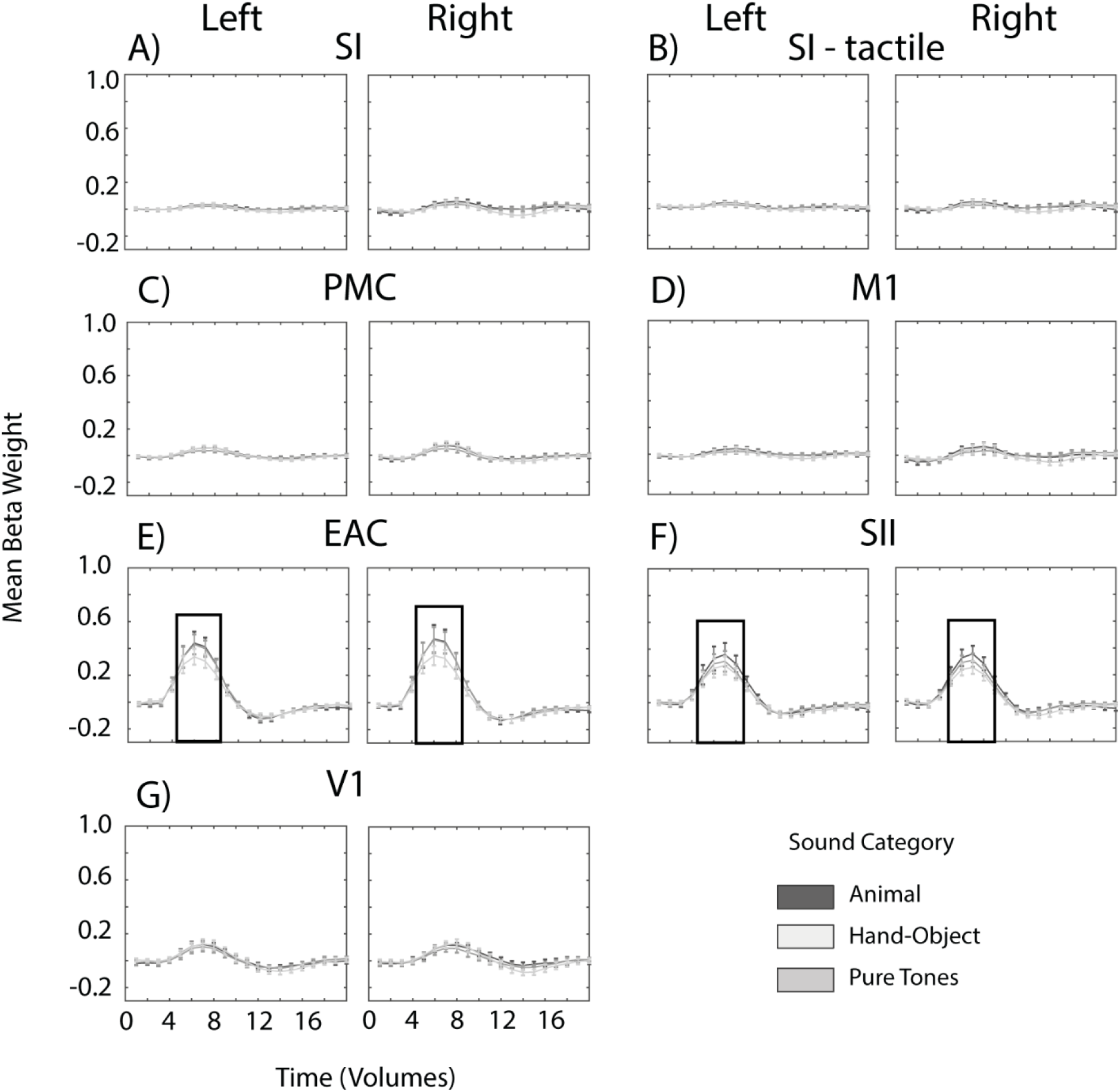
Univariate Analysis. (A) Mean Beta Weight plotted for the SI ROI for each sound category and split by hemisphere. (B) As in A but for the top 100 voxels that were responsive to tactile stimulation of the hands in an independent localizer session. (C-G) as in A but for the remaining ROIs. The rectangle in figures E&F highlight where the data was extracted around the peak (5-8s) for statistical analysis.

## Discussion

In the present study we show that hearing sounds depicting familiar hand-object interactions elicit significantly different patterns of activity in primary somatosensory cortex (SI), despite the complete absence of external tactile stimulation. Such decoding was not found for two control categories of familiar animal vocalizations, and unfamiliar pure tones, in any analysis involving SI. Moreover, decoding accuracies were significantly higher for hand-object interaction sounds compared to pure tones when restricted to the independently localized hand-sensitive voxels of the left SI (or when pooling across hemispheres using the entire SI). We further show that SII – a higher order somatosensory region – shows reliable decoding for all sound categories presented. Control analyses demonstrated that these effects in SI and SII were not driven by proximity to nearby regions also displaying significant effects (PMC and EAC respectively). Thus, fine-grained information about different sound categories is present within both primary (SI – hand-object interactions) and higher somatosensory brain areas (SII - hand-object interactions, animal vocalizations and pure tones).

### Representation of Sound Categories in Somatosensory Cortices

The present findings are in broad agreement with previous work showing auditory responses in early somatosensory areas (Zhou and Fuster 2004; Beauchamp and Ro 2008; Lemus et al. 2010; Pérez-Bellido et al. 2018). In particular, the current study partially replicates Pérez-Bellido et al. (Pérez-Bellido et al. 2018) by showing reliable discrimination of pure tone stimuli in SII, even after eroding voxels close to the border with early auditory areas to control the possible influence of partial volume effects. The current study extends this previous work by showing that A) SII further discriminates different exemplars within specific high level sound categories (hand-object interactions and animal vocalizations) and that B) SI contains representations that discriminate particular high level sound categories (hand-object interactions) but not others (animal vocalizations and pure tones) and may manifest a degree of functional specialization for specific sound categories (Left SI - tactile sensitive voxels and also in Pooled SI entire region for hand-object interaction sounds vs pure tones).

Interestingly, when analyses were split by hemisphere and by different putative areas (specifically SI) in Pérez-Bellido et al. (Pérez-Bellido et al. 2018), effects were only found when voxels responsive to auditory stimulation in an independent localizer were used to define the regions. Defining the regions using voxels responsive to tactile stimulation did not lead to any reliable effects for their pure tone stimuli in putative SI. In contrast, in the present experiment we show that selecting voxels responsive to tactile stimulation of the hand produced robust discrimination in Left SI for sounds depicting hand-object interactions but not others (animal vocalizations or pure tones). Thus, it might be the case that early somatosensory regions contain information about multiple types of sound category but that the method used to select voxels for input to the analysis is critical in determining the effects that are found. This may be particularly the case given that somatosensory cortices are involved in more than just sensory processing of touch – being also implicated in processing of emotional states for instance (Adolphs 2002; Keysers et al. 2010).

The findings observed in SII (and also premotor cortex – see below) of reliable decoding for all sound categories suggest the involvement of the dorsal auditory stream (Rauschecker and Tian 2000; Bizley and Cohen 2013). This pathway - originating in posterior auditory regions and linking to IPL and then premotor areas - is thought to be involved in linking sounds to the actions that have produced them and as such in sensorimotor integration and control (Rauschecker 2012). Importantly neurons responsive to tactile stimulation are present in posterior auditory cortex (e.g. Fu et al. 2003; see also Rauschecker and Scott 2009) and these may contribute towards the effects we observe in SII (see Pérez-Bellido et al. 2018).

### Triggering cross-modal content-specific information in SI from audition

The present study agrees with a set of studies that show fine-grained representation of distal sensory categories even in the earliest regions of supposedly unimodal primary sensory areas, across multiple sensory domains (Meyer et al. 2010; Vetter et al. 2014; Smith and Goodale 2015; see also Pérez-Bellido et al. 2018). Our results significantly extend this previous body of work by demonstrating, for the first time, that a similar effect is present across the domains of audition and touch for particular familiar sound categories. It is important to note, furthermore, that the pattern of decoding performance was different in SI and in auditory cortex, with better decoding of pure tones than hand-object interactions in auditory cortex whereas SI showed some evidence for better decoding of hand-object interactions than pure tones in particular analyses (Left SI tactile sensitive voxels, SI entire region pooled across hemispheres). As such we suggest that the present results in SI are unlikely to reflect passive relay of low-level acoustic features from auditory cortex via the auditory dorsal stream (Rauschecker and Tian 2000; Bizley and Cohen 2013).

The current results expand on our earlier study that showed simply viewing images of familiar graspable objects led to reliable decoding in SI (Smith and Goodale 2015; see also Meyer et al. 2011). Thus either viewing images or hearing sounds of particular objects – i.e. those related to hand-object interactions-triggers fine-grained representations of those objects in SI. What might be the function of this cross-sensory information being present in SI? If Predictive Processing is assumed to be the general computational function of the brain (Friston 2009; Friston et al. 2009; Clark 2013), then it may be the case that either seeing a familiar graspable object, or hearing the sounds depicting a familiar hand-object interaction, leads to fine-grained activation of associated tactile/haptic features in SI that is useful for future interaction with the specific object (see Pérez-Bellido et al. 2018 for a related explanation; see also Jacobs and Xu 2019). The strength of this effect would be expected to be determined by the magnitude of co-occurrence of sensory stimulation across audition and touch for particular object categories (Meyer and Damasio 2009; see also Jacobs and Xu 2019).

How does auditory stimulation lead to content-specific information being present in SI? There are several possible anatomical routes through which this could be accomplished (see e.g. Driver and Noesselt 2008). First, auditory information may first arrive at high level multisensory convergence zones, such as pSTS, posterior parietal cortex or ventral premotor areas, before such areas then send feedback to SI (see Driver and Noesselt 2008; Meyer et al. 2009; see also Smith and Goodale 2015). Another possible high-level convergence site is the fusiform gyrus, since Kassuba et al. (Kassuba et al. 2013) found semantically coherent auditory and haptic object features activated this area. Second, auditory information may be directly projected to SI, without passing through such higher multisensory regions. Such direct connections have been found in animal models between certain sensory pairings: e.g. from primary auditory to primary visual cortex (e.g. Falchier et al. 2002, 2010) and between primary auditory and somatosensory cortex (Budinger et al. 2006; see also Cappe and Barone 2005). However it has been proposed that such direct connections are relatively sparse as opposed to the amount of feedback arriving from higher multisensory areas (Driver and Noesselt 2008). Finally a third possibility is the involvement of lower tier multisensory regions (Driver and Noesselt 2008) that are anatomically located next to primary sensory areas: for instance auditory regions located close to SII may be bimodal responding to both auditory and tactile information (see e.g. Wallace et al. 2004; Cappe and Barone 2005). Our current results clearly highlight robust information about all sound categories in both SII and also in premotor cortices (Right – all categories; Left – hand object interactions and animal vocalizations) – and as such these regions may be involved in transmitting information back to SI about specific sound categories. However, this would not explain why decoding in Left SI (hand sensitive voxels) is higher for hand-object interactions than pure tones – the opposite of the pattern observed in SII, while no differences between categories were observed in premotor areas. Thus, the specific route by which such information is transmitted to SI is something that will need to be addressed in future research.

### The role of prior audio-haptic experience

In the present study we only ever found reliable decoding of hand-object interactions in SI regardless of whether the analysis used the entire SI anatomical mask or a small subset of tactile sensitive voxels. Critically, when limited to hand sensitive voxels in Left SI was there a reliable difference between hand-object interactions and our low-level pure tone control stimuli. This difference was also apparent when pooling across hemispheres using the SI anatomical mask. Thus, we demonstrate a degree of functional specialization for hand-object interactions in some SI analyses: particularly in the Left SI hand sensitive voxels. However, it is important to highlight that we did not find any evidence of significantly greater decoding for hand-object interaction sounds than our high-level control of animal vocalizations. As such, it is possible that similar effects may be present in SI for familiar sound categories in general, especially when one considers the multifaceted role of SI in both somatosensation but also higher-level socio-emotional processing (Keysers et al. 2010). Future studies should set out to explicitly dissociate the influence of these features on cross-modal activity in early somatosensory regions.

### Decoding Sound Categories in (Pre-) Motor Regions

In Left PMC we found reliable decoding of both familiar categories whereas in the Right and Pooled PMC we found reliable decoding of all sound categories. We did not find any differences between sound categories in any PMC sub-region. Thus, premotor cortex contains information about all sound categories presented, across the two hemispheres. Finding reliable decoding for multiple sound categories in PMC is consistent with multiple theoretical accounts: first, the relatively non-specific pattern of decoding implicates the auditory dorsal stream (Rauschecker and Tian 2000; Bizley and Cohen 2013) which terminates in premotor areas and would be expected to contain information about all types of sound presented. While PMC is also known to play a large role in processing action related information through a potential mirroring mechanism (Gallese et al. 1996; Gazzola et al. 2006), this account would predict greater decoding for sounds strongly implying an association action (e.g. hand-object interaction & animal vocalizations) which we did not find in the current data (although Left PMC only showed decoding for both familiar object categories but not pure tones – which may be partially in agreement with this view). Finally, the absence of decoding in primary motor areas (M1) is broadly consistent with past work in this area (Lewis et al. 2005; Smith and Goodale 2015).

## Conclusion

We have shown that the identity of familiar hand-object interaction sounds can be discriminated in SI, in the absence of any concurrent tactile stimulation. Our work provides converging evidence that activity in supposedly modality-specific primary sensory areas can be shaped in fine-grained manner by relevant contextual information originating in different sensory modalities (Meyer et al. 2010, 2011; Vetter et al. 2014; Smith and Goodale 2015). Such an effect implies the potential for fine-grained multisensory interactions even in the earliest regions of sensory cortex.

