## Supplementary Material for "Decoding sounds depicting hand-object interactions in primary somatosensory cortex"

**Supplementary Text**

***Similarity Rating Experiment***

***Participants***

23 participants (22 females; one male) took part in the similarity rating experiment with an average age of 19. Participants were all right-handed, Undergraduate Psychology students at the University of East Anglia (UEA), with normal, or corrected-to-normal hearing. Participants were recruited on a voluntary basis using the School of Psychology Research Participation System (SONA), and were awarded four credits in exchange for their participation.

***Stimuli and Design***

All 30 sounds selected for use in the main experiment were used in this task. Each sound was paired with every other possible sound, including those of the same category, resulting in 435 total similarity rating trials (each pairwise combination of 30 sounds). Sounds were presented in a randomized order for each participant in EPRIME software. The experiment was presented to participants on a 1920x1080 HP computer screen, and they heard the trials through a pair of SONY 7506 headphones which were set to a self-reported comfortable volume level. Participants were told to rate the similarity of each pair of sounds played on a 7 point likert scale (1=no similarity, 7=extremely similar).

**Results**

We extracted the average within category similarity between each pair of sounds from the same category (excluding the two exemplars of the same sound – e.g. two examples of knocking on a door) and compared these across different categories (hand object interactions, animal vocalizations and pure tones – see Supp. Figure 1). The analysis revealed that mean similarity within categories was similar for hand object and animal sounds ($M = 1.75$ and 1.7, respectively – $p = .424$; two tailed signed rank test) both of which were significantly lower than that for the pure tones ($M = 5.08$; both p's $<.0001$; two tailed signed rank test). We also examined whether the average similarity of the two examples of the same sound differed across categories (i.e. the two different examples of a dog barking – the main diagonal of Supp. Figure 1). Again we found that there was no significant difference between hand object interactions and animal vocalizations ($M = 5.6$ and 5.3 respectively, $p = .098$) but both these categories were reliably different from pure tones ($M = 6.78$; both p's $< .0001$; two tailed sign rank test).

Thus the similarity across different exemplars within a category is approximately equal across the two familiar categories used in the present experiment (Hand object interactions and animal vocalizations) both of which are noticeably smaller than the similarity between different exemplars within the pure tone category. Given the simple and artificial nature of the pure tone control stimuli (simple differences in frequency) these differences with more ecologically relevant categories are not unexpected.

### Sound Selection Pilot Rating Experiment

A pilot behavioural experiment was designed to determine which sounds would be used in the main experiment. Participants listened to a selection of different sounds, and were asked to identify them, and rate them on a number of different aspects.

### Pilot Experiment – Methods

#### *Participants*

Psychology undergraduate students (*N* = 29; 5 male) were recruited for this experiment, with an age range of 18-37 years (*M* = 20.36, *SD* = 3.42). All participants reported normal or corrected-to-normal vision, and normal hearing. Participants signed written consent following ethical approval from the Research Ethics Committee of the School of Psychology at the University of East Anglia. Participants were reimbursed with four SONA credits for their time, in line with the UEA SONA credit payment guidelines.

#### *Stimuli and Design*

Initially, three sound categories were piloted: Hand-object interactions (e.g. typing on a keyboard, knocking on a door), mouth-object interactions (e.g. eating an apple, sipping a drink), and animal vocalizations (e.g. dog barking, rooster crowing). Royalty free sounds in WAV format were downloaded from various sound databases such as Soundsnap.com, YouTube.com, and from a sound database used in Giordano, McDonnel, and McAdams (Giordano et al. 2010). Using Audacity audio software 2.1.2, all sounds were cut to exactly 2000ms in length, ensuring sound filled the entire duration. Sounds were all normalised to the root mean square (Giordano et al. 2013). Overall, 66 different stimuli were piloted; 33 per category, with two exemplars of each stimulus. The experimental session lasted between 45-60 minutes for each participant.

For the experimental task, sounds were loaded into an E-Prime 2.0 experiment. Participants listened to each sound once through professional SONY MDR-7506 headphones, with the volume set at a self-reported comfortable level (as in Leaver and Rauschecker 2010;

Meyer et al. 2010; Man et al. 2012, 2015). To begin a trial, participants were asked to press a button, which would initiate a countdown screen from three seconds. Following the countdown, a 2000ms sound was played whilst participants viewed a blank white screen. All sound stimuli were presented in a random order. Once a sound finished playing, participants were automatically redirected to a screen asking them a series of self-paced questions. The questions used in this experiment were derived from a series of previous research using sounds (Marcell et al. 2000; Schneider et al. 2008; Giordano et al. 2010, 2013). The following seven questions were asked:

(1) *Identification*. Participants were asked to identify the sound using at least one verb, and one or two nouns, as seen in Giordano et al. (Giordano et al. 2010). Participants were instructed to make their best guess if they did not know.

(2) *Confidence*. Participants were asked to rate how confident they were with their decision. Ratings we on a 1-7 Likert scale from 1 (not at all confident) to 7 (very confident).

(3) *Familiarity*. Participants rated how familiar they were with the sound. In particular, how commonly they heard the sound in day-to-day life, not just in the way it was presented to them. This was important, since the main study was interested in how general familiarity with a sound may evoke traces of activity in other brain areas. Ratings were made from 1 (not at all familiar) to 7 (very familiar).

(4) *Number of sound-generating events.* The next rating was how many sound-generating events participants believed were present. For example, a ticking of a clock would have many events, whereas a simple click of a mouse button would only have one event. It was important to control for this across our sound categories, since the number of sound-generating events has been found to evoke different brain activity patterns important for classifier performance (Meyer et al. 2011). Ratings were made from 1 (no events) to 7 (many events).

(5) *Action and movement related information*. Next, participants were asked to subjectively rate the amount of action and movement related information that was present for each sound – whilst this was expected to be higher for hand- and mouth- object interactions, participants were given no indication to this. They were simply asked “Did the sound convey action and movement related information?” Ratings were made from 1 (no action and movement related information) to 7 (much action and movement related information).

(6) *Vividness*. Participants also rated how strongly they experienced mental imagery whilst listening to the sound, with the question adapted from the Bucknell Auditory Imagery Scale (Halpern 2015), and also Meyer et al. (Meyer et al. 2010). Participants were asked to rate

“the quality of the sound in terms of how strongly it evoked an image in your head”. Ratings were made from 1 (no image evoked at all) to 7 (I could see the image very clearly).

(7) *Perspective.* Finally, participants were asked to specify the perspective they imagined the sound to be taking place. Participants were given five options: 1. You were making the sound yourself. 2. Somebody else was making the sound. 3. A (non-human) animal was making the sound. 4. Nobody was making the sound. 5. Other (please specify).

### Results

Stimuli to be used in the main experiment were primarily selected according to correct identification (at least 90% across all participants), with high confidence and familiarity ratings (> 5). Identification was analysed as strict correct (correct verb and noun, e.g. door knock) or a not so strict correct (either a verb or a noun, e.g. knocking). We also matched the average number of sound-generating events across our final sound categories (see Supp. Table 1 for final stimuli ratings).

For the final stimulus set, it was decided that mouth-object sounds would be removed due to the ambiguity of these sounds conveying purely a mouth-related action. For example, sounds such as eating an apple or brushing teeth would also involve a hand movement. Thus, ratings from the mouth-object sounds have been excluded. Following this decision, we then decided to include pure tones as an unfamiliar control category. Once the final stimulus set was decided, sounds were re-normalised to the root mean square (Giordano et al. 2013).

#### *Final selected stimuli:*

1) Familiar sounds depicting hand-object interactions: Five sub-categories; typing on a keyboard, bouncing a basketball, knocking on a door, crushing paper, and sawing wood.
2) Familiar control stimuli: Five animal vocalizations; dog barking, birds chirping, rooster crowing, fly buzzing, and frog croaking. Animal sounds were chosen as a familiar sound control category, to determine whether any familiar sound can evoke traces of activity to primary somatosensory cortex (Lewis, 2005; Lewis, Phinney, Brefczynski-Lewis, & DeYoe, 2006; Lewis, Talkington, Puce, Engel, & Frum, 2011).
3) Unfamiliar control stimuli: Five pure tones (different frequencies of the same tone; 400Hz, 800Hz, 1600Hz, 3200Hz, and 6400Hz). Tones were included as an unfamiliar control category (Lewis, 2005; Mesulam, 1998) to rule out the idea that merely any sound can lead to discrimination in primary somatosensory cortex.

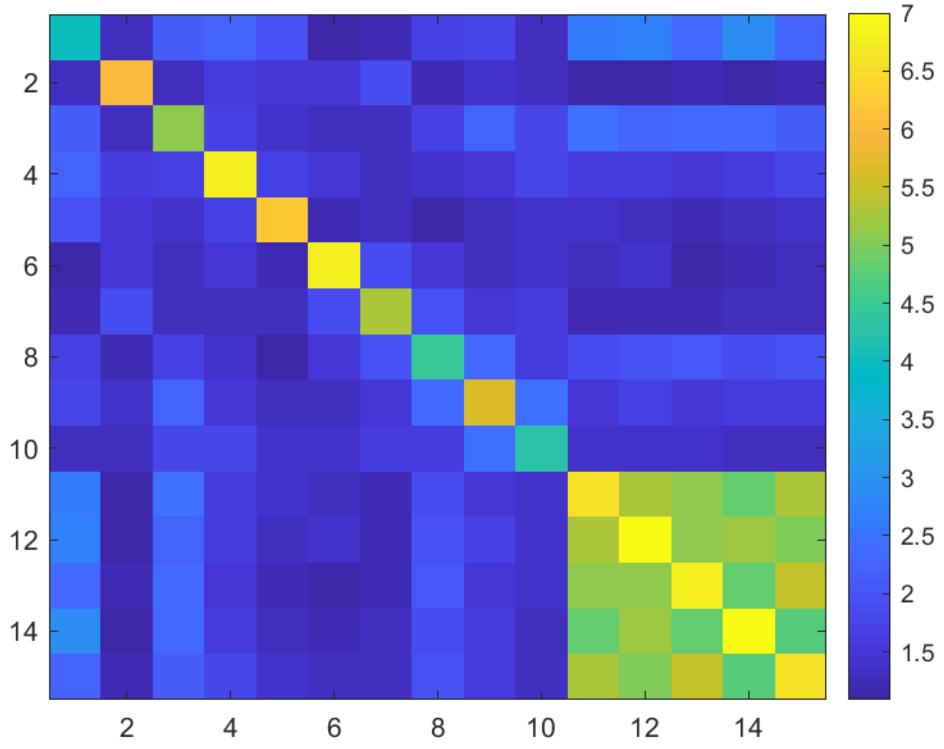

**Supplementary Figure 1**: Perceived similarity between all sounds used in the main experiment (higher scores = greater similarity) rated by an independent group of participants (N=23). The matrix is organized such that the first 5 rows (& columns) correspond to animal vocalizations, the next 5 to hand-object interactions and the final 5 to pure tone stimuli. The main diagonal is not zero as this position indicates the similarity of the two different versions of each sound to one another (e.g. the similarity between two examples of typing on a keyboard).

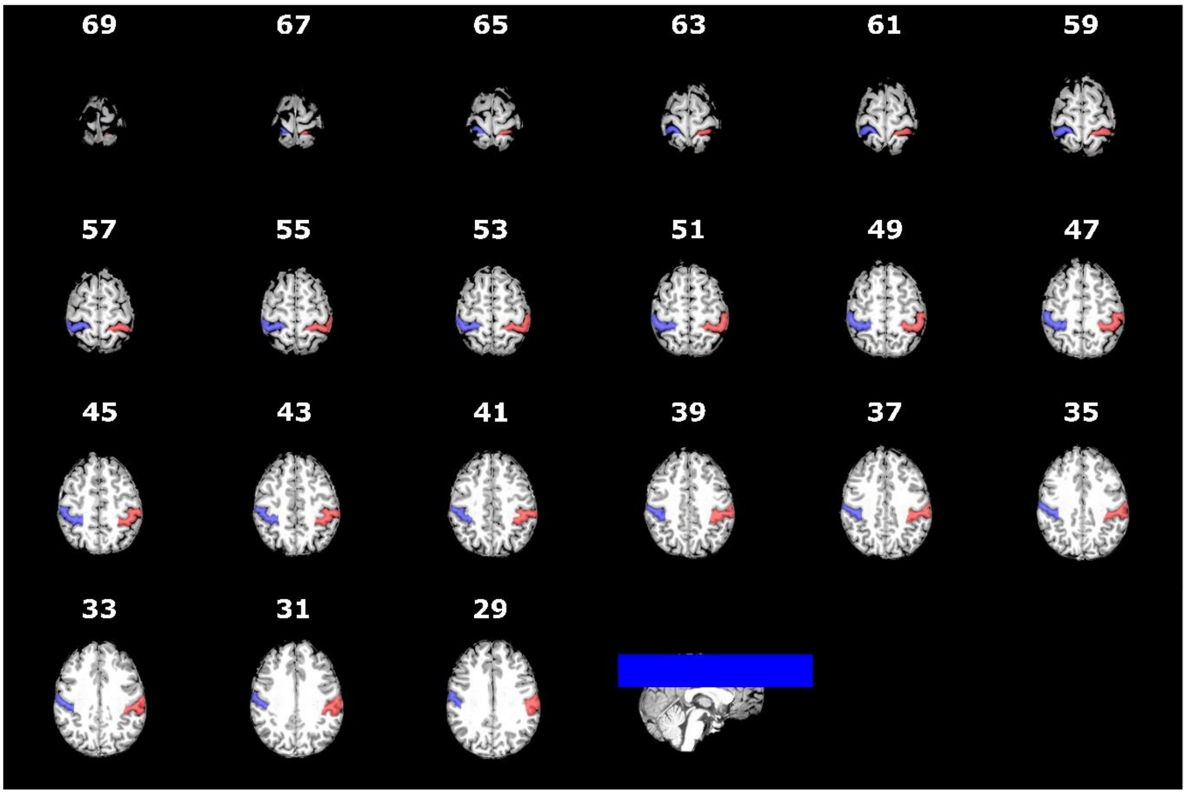

**Supplementary Figure 2**: Hand-drawn anatomical masks of the lateral post-central gyrus for a representative participant (the same as in Figure 1 of the main manuscript). The numbers in white refer to slices through the Z plane. The box in the lower right image depicts the slices of the brain on which the PCG was marked (see Methods).

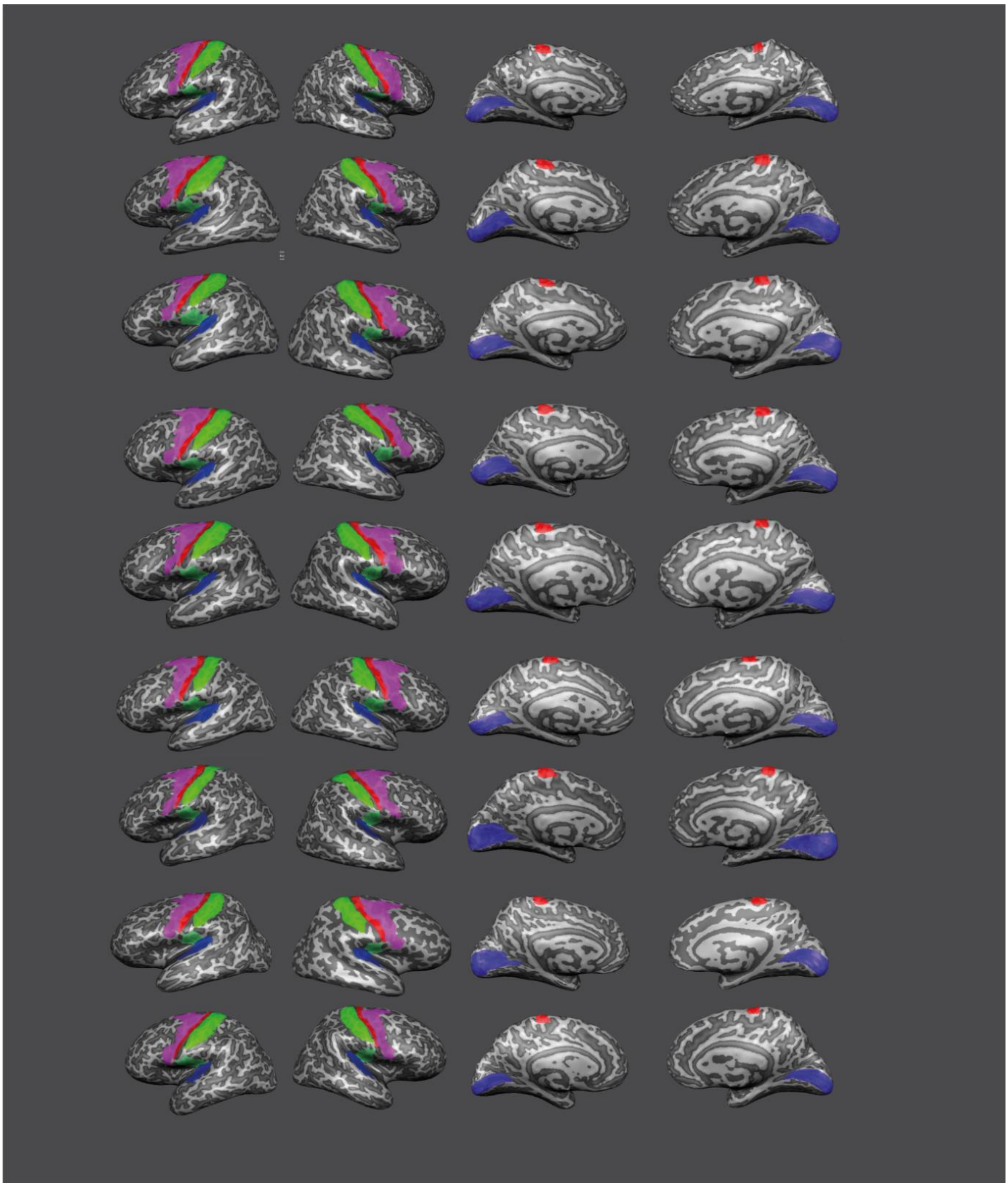

**Supplementary Figure 3**: Regions of Interest (ROIs) defined per participant using a hand-drawn mask (SI – bright green) or the Glasser parcellation (SII - green, M1 - red, PMC - pink, EAC – dark blue, V1 – light blue) for all participants except the participant shown in Figure 1. Each ROI is shown on inflated cortical surface reconstructions of both Left and Right hemispheres (both lateral and medial views are shown).

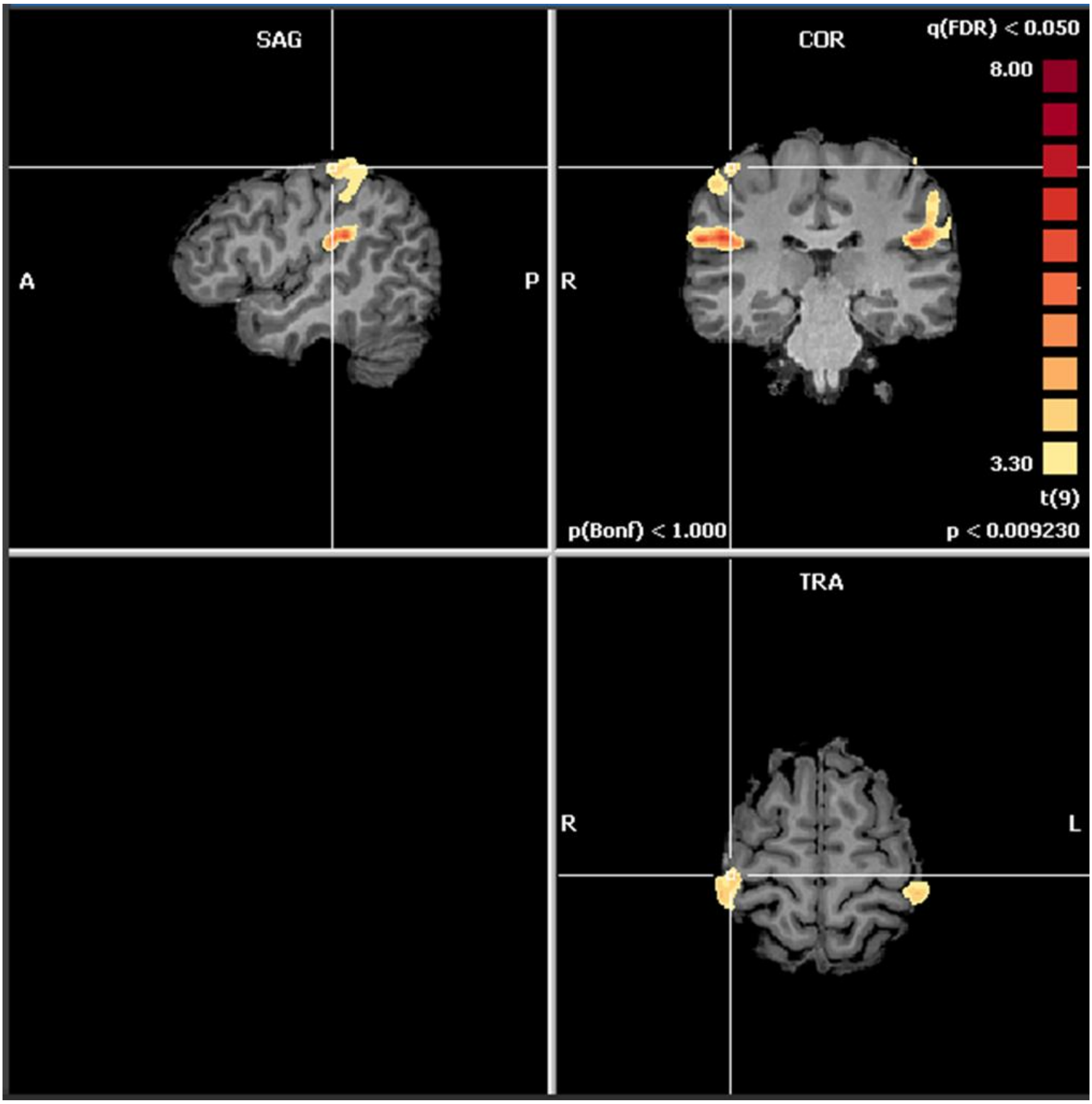

**Supplementary Figure 4:** Group activation of the somatosensory functional localizer. Displayed voxels are significant at the group level (RFX, $t > 3.3$, $p < .01$, uncorrected; voxel extent threshold = 50) for the contrast of tactile stimulation of the hands vs baseline. As expected, robust activation was found over both SI and SII. See Figure 1 for an example of individual participant activation in this task.

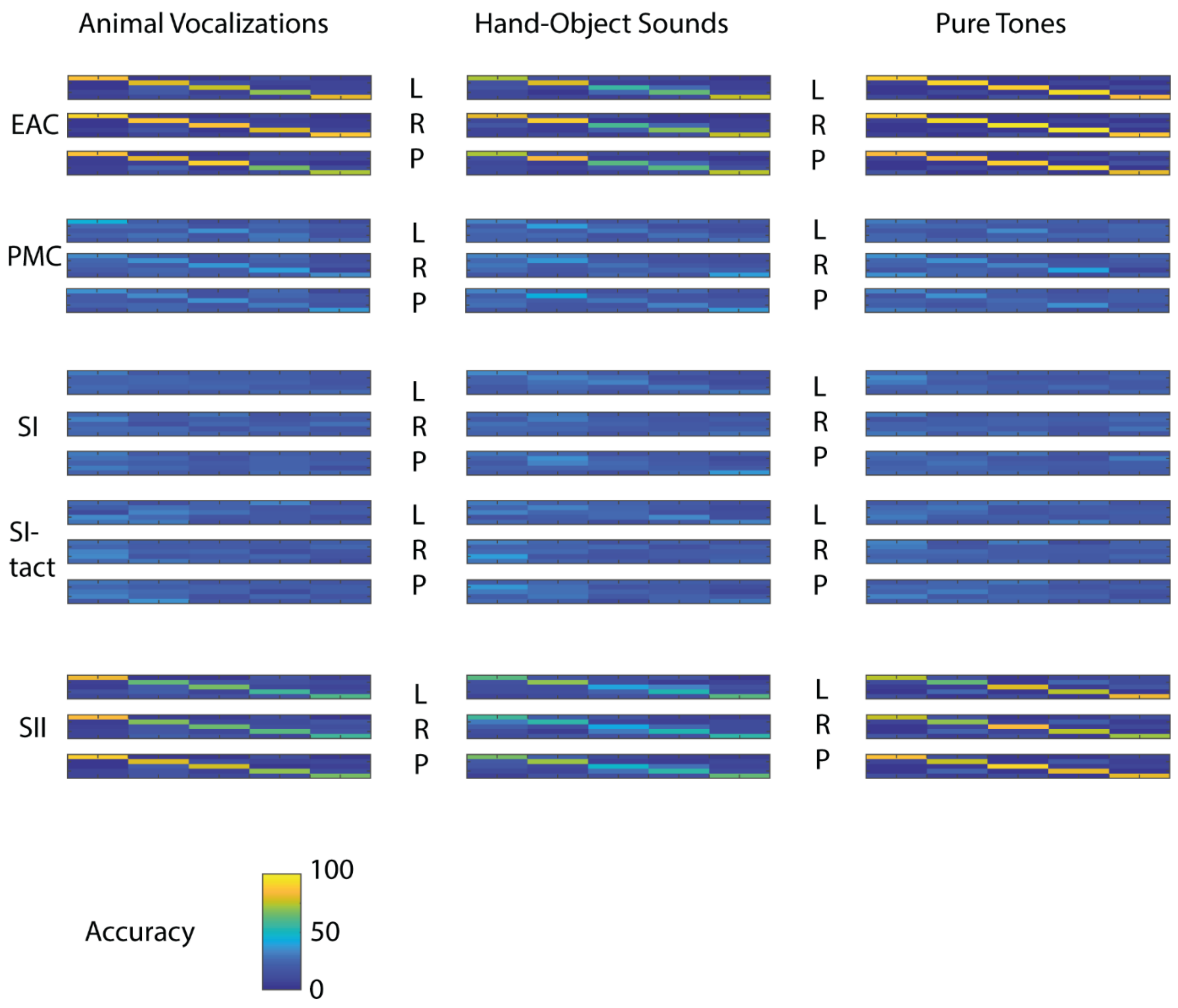

**Supplementary Figure 5**: Confusion Matrices underlying classification performance for each ROI displaying significant decoding results. Panel columns represent the different sound categories (animal vocalizations, hand-object interactions and pure tones) whereas rows represent each different ROI split by Left, Right or Pooled across hemisphere. Each 5X5 matrix specifies for a given ROI and sound category, the specific exemplar presented (row) and the specific exemplar chosen by the classifier (column). As such the diagonal represents correct classification performance while the off-diagonal represents errors.

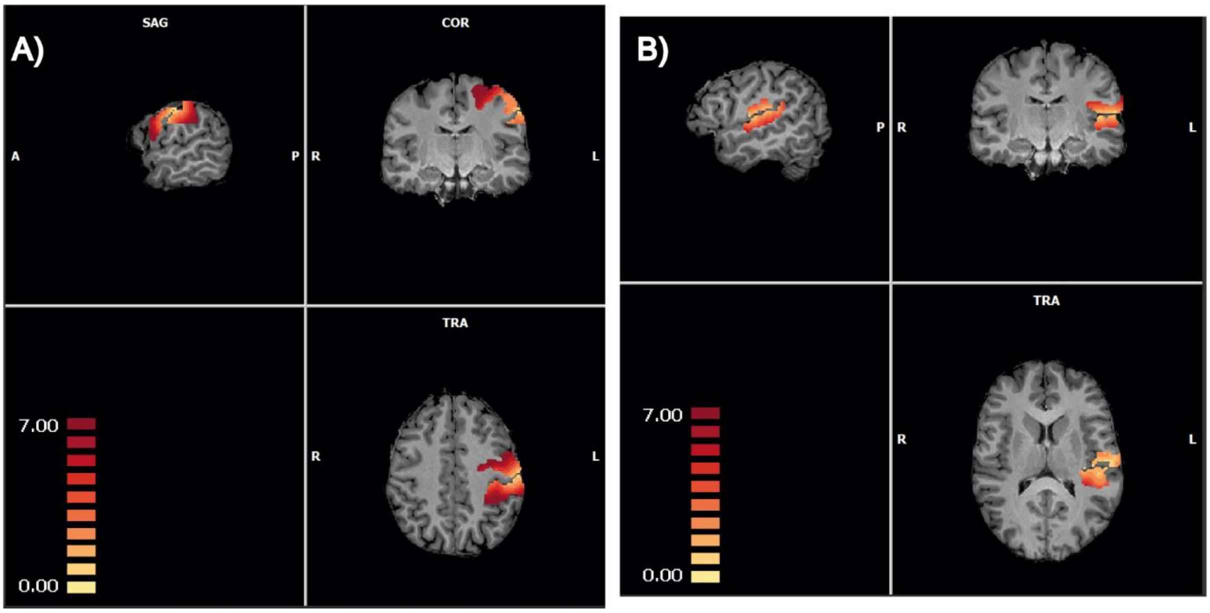

**Supplementary Figure 6**: (A) Volume map depicting the different levels of erosion used for eroding the nearest voxels between PMC and SI. (B) as in A except for EAC and SII. The colour scale represents the voxel erosion level assigned to each voxel (1-7 mm; warmer colours equal more distant).

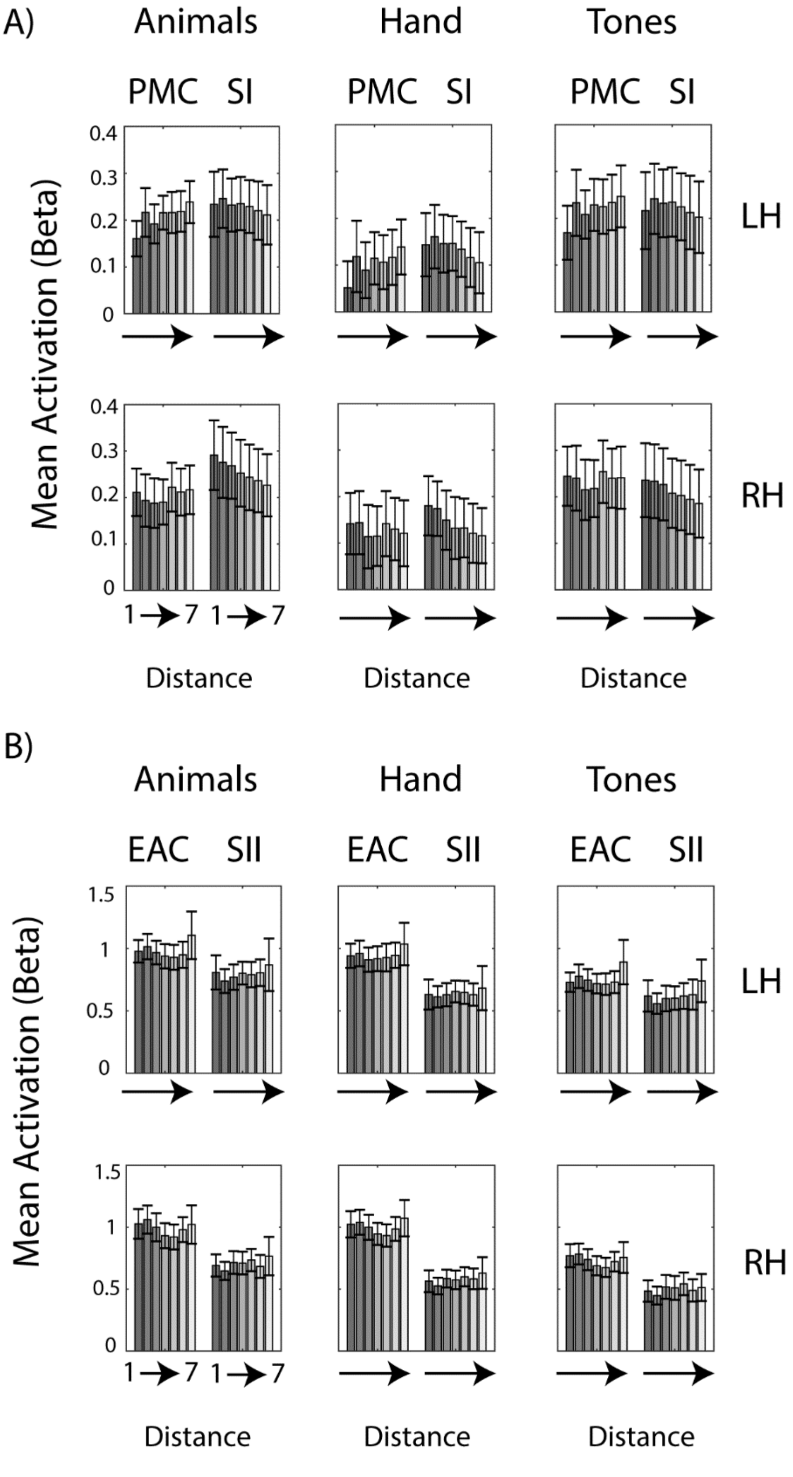

**Supplementary Figure 7**: (A) Univariate signal in PMC and SI for each sound category and each ROI (split by hemisphere - rows), as a function of the level of voxel erosion employed to eliminate the contribution of nearby voxels. (B) As in A except for EAC and SII.

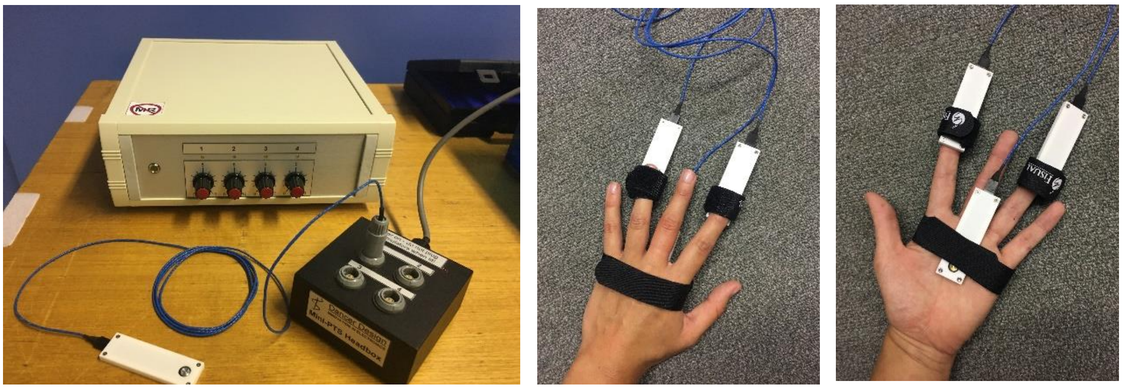

**Supplementary Figure 8*:*** Image of miniature Piezo-Tactile Stimulator. Demonstration of three pads placed on the index finger, ring finger, and palm of the left hand.

| *Stimuli* | *Confidence* | *Familiarity* | *Sound gen events* | *Action info* | *Perspective* | *Vividness* | *Strict correct* | *Not strict correct* |
|---|---|---|---|---|---|---|---|---|
| *Typing keyboard 1* | 5.93 | 5.89 | 5.68 | 5.07 | 1.75 | 5.68 | 64.29% | 96.43% |
| *Typing keyboard 2* | 5.57 | 5.86 | 5.43 | 5.11 | 1.54 | 5.43 | 60.71% | 89.29% |
| *Door knock 1* | 6.79 | 6.25 | 5.46 | 5.50 | 1.79 | 5.93 | 96.43% | 100.00% |
| *Door knock 2* | 6.36 | 5.86 | 5.07 | 5.43 | 1.75 | 5.89 | 82.14% | 85.71% |
| *Sawing wood 1* | 6.36 | 4.71 | 3.93 | 5.50 | 1.82 | 5.79 | 67.86% | 100.00% |
| *Sawing wood 2* | 5.79 | 4.36 | 4.64 | 5.61 | 1.89 | 5.93 | 67.86% | 92.86% |
| *Basketball bounce 1* | 5.79 | 4.75 | 4.82 | 5.39 | 1.86 | 6.07 | 78.57% | 85.71% |
| *Basketball bounce 2* | 6.32 | 4.82 | 4.14 | 5.50 | 2.14 | 5.89 | 78.57% | 85.71% |
| *Paper crush 1* | 5.54 | 5.32 | 3.54 | 5.11 | 2.04 | 5.00 | 64.29% | 100.00% |
| *Paper crush 2* | 4.93 | 5.32 | 3.96 | 5.11 | 1.79 | 5.36 | 92.86% | 96.43% |
| ***AVERAGE:*** | **5.94** | **5.31** | **4.67** | **5.33** | **1.84** | **5.70** | **75.36%** | **93.21%** |
| *Dog bark 1* | 6.89 | 6.07 | 4.68 | 4.00 | 2.96 | 6.43 | 100.00% | 100.00% |
| *Dog bark 2* | 6.93 | 6.07 | 4.75 | 3.96 | 3.00 | 6.25 | 96.43% | 100.00% |
| *Bird chirp 1* | 6.75 | 6.46 | 5.29 | 4.04 | 3.00 | 6.04 | 85.71% | 100.00% |
| *Bird chirp 2* | 5.39 | 4.89 | 4.96 | 3.61 | 3.00 | 5.04 | 71.43% | 96.43% |
| *Rooster crow 1* | 6.54 | 4.61 | 3.82 | 3.00 | 3.07 | 5.86 | 57.14% | 100.00% |
| *Rooster crow 2* | 6.50 | 4.79 | 3.75 | 3.29 | 3.00 | 5.89 | 60.71% | 100.00% |
| *Frog croak 1* | 6.57 | 4.43 | 4.21 | 3.32 | 3.00 | 5.43 | 60.71% | 100.00% |
| *Frog croak 2* | 6.61 | 4.61 | 4.07 | 3.32 | 2.93 | 5.86 | 60.71% | 100.00% |
| *Fly buzz 1* | 6.11 | 5.54 | 4.39 | 5.18 | 2.96 | 5.82 | 67.86% | 96.43% |
| *Fly buzz 2* | 6.32 | 5.64 | 4.96 | 4.89 | 3.00 | 6.00 | 71.43% | 100.00% |
| ***AVERAGE:*** | **6.46** | **5.31** | **4.49** | **3.86** | **2.99** | **5.86** | **73.21%** | **99.29%** |

**Supplementary Table 1: Mean ratings of selected stimuli (N=29)**

| Region | Mean | Standard Deviation |
|---|---|---|
| LEAC | 7078 | 828 |
| LM1 | 7846 | 1011 |
| LPMC | 19042 | 1832 |
| LSI | 16848 | 1717 |
| LSII | 4910 | 854 |
| LV1 | 10833 | 3381 |
| REAC | 6183 | 820 |
| RM1 | 7758 | 728 |
| RPMC | 19300 | 2205 |
| RSI | 15750 | 1830 |
| RSII | 4648 | 809 |
| RV1 | 11517 | 3446 |

**Supplementary Table 2: Mean volume of each ROI across participants (in cubic mm)**
EAC=Early Auditory Cortex (see Methods), PMC = Pre-motor cortex.
